# Bivalent mRNA vaccine improves antibody-mediated neutralization of many SARS-CoV-2 Omicron lineage variants

**DOI:** 10.1101/2023.01.08.523127

**Authors:** Nannan Jiang, Li Wang, Masato Hatta, Chenchen Feng, Michael Currier, Xudong Lin, Jaber Hossain, Dan Cui, Brian R. Mann, Nicholas A. Kovacs, Wei Wang, Ginger Atteberry, Malania Wilson, Reina Chau, Kristine A. Lacek, Clinton R. Paden, Norman Hassell, Benjamin Rambo-Martin, John R. Barnes, Rebecca J. Kondor, Wesley H. Self, Jillian P. Rhoads, Adrienne Baughman, James D. Chappell, Nathan I. Shapiro, Kevin W. Gibbs, David N. Hager, Adam S. Lauring, Diya Surie, Meredith L. McMorrow, Natalie J. Thornburg, David E. Wentworth, Bin Zhou

**Affiliations:** COVID-19 Emergency Response, Centers for Disease Control and Prevention, Atlanta, GA, USA; Influenza Division, National Center for Immunization and Respiratory Diseases, Centers for Disease Control and Prevention, Atlanta, Georgia, USA; Division of Viral Diseases, National Center for Immunization and Respiratory Diseases, Centers for Disease Control and Prevention, Atlanta, Georgia, USA; Vanderbilt University Medical Center, Nashville, TN, USA; Beth Israel Deaconess Medical Center, Harvard University, Cambridge, MA, USA; Wake Forest Baptist Medical Center, Winston-Salem, NC, USA; Johns Hopkins University, Baltimore, MD, USA; University of Michigan, Ann Arbor, MI, USA

## Abstract

The early Omicron lineage variants evolved and gave rise to diverging lineages that fueled the COVID-19 pandemic in 2022. Bivalent mRNA vaccines, designed to broaden protection against circulating and future variants, were authorized by the U.S. Food and Drug Administration (FDA) in August 2022 and recommended by the U.S. Centers for Disease Control and Prevention (CDC) in September 2022. The impact of bivalent vaccination on eliciting neutralizing antibodies against homologous BA.4/BA.5 viruses as well as emerging heterologous viruses needs to be analyzed. In this study, we analyze the neutralizing activity of sera collected after a third dose of vaccination (2-6 weeks post monovalent booster) or a fourth dose of vaccination (2-7 weeks post bivalent booster) against 10 predominant/recent Omicron lineage viruses including BA.1, BA.2, BA.5, BA.2.75, BA.2.75.2, BN.1, BQ.1, BQ.1.1, XBB, and XBB.1. The bivalent booster vaccination enhanced neutralizing antibody titers against all Omicron lineage viruses tested, including a 10-fold increase in neutralization of BQ.1 and BQ.1.1 viruses that predominated in the U.S. during the last two months of 2022. Overall, the data indicate the bivalent vaccine booster strengthens protection against Omicron lineage variants that evolved from BA.5 and BA.2 progenitors.

## Introduction

Hundreds of SARS-CoV-2 variant lineages have emerged, spread, and replaced by newer lineages in the past three years. Thirteen of them were designated as “variants of concern” (VOCs) or “variants of interest” (VOIs) by the World Health Organization (WHO) with Greek alphabet letters assigned from Alpha to Omicron. Each of these so-called “variants” exists as a lineage that includes many descendant lineages (a.k.a. sublineages or clades) defined by specific genetic changes in the genome (e.g., Pango Nomenclature^1^). In the United States, viruses containing primarily a D614G substitution in the spike protein predominated in 2020 followed by the Alpha lineage viruses predominating in the first half of 2021 and the Delta lineage viruses in the second half of 2021^2^. The early Omicron lineage, BA.1, rapidly displaced the Delta lineage between December 2021 and January 2022 (Fig. 1), with a fitness advantage largely conferred by immune evasion^2, 3, 4, 5, 6, 7, 8^. Omicron viruses evolved further leading to several fit lineages, many of which accumulated additional spike substitutions in the course of infecting or reinfecting people. In 2022, BA.1 and its descendant, BA.1.1, predominated in January-March; BA.2 and its descendant, BA.2.12.1, predominated in March-June; and BA.5 predominated in July-October until BQ.1 and BQ.1.1 surpassed it in November-December (Fig. 1).

**Fig. 1:**
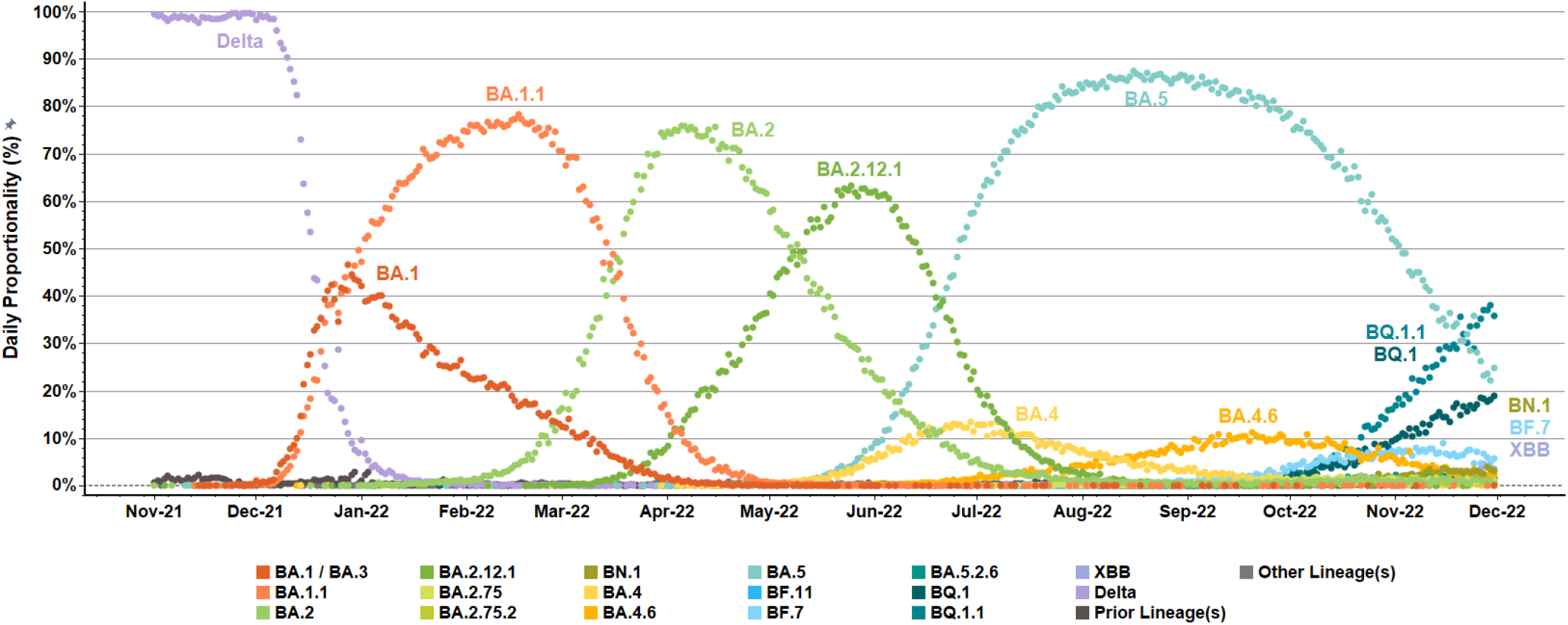
Emergence and diversification of SARS-CoV-2 Omicron variants and lineages in US surveillance networks. Daily, percent proportionalities (dot icons, 0% to 100%) were summarized for detected variant/lineage populations within US National SARS-CoV-2 Strain Surveillance (NS3) and baseline surveillance initiatives from November 1, 2021, to November 30, 2022. Reported sequences were aggregated and color-coded based on attributed Pango lineages. Delta (B.1.617.2) variant preceded the emergence and expansion of Omicron (B.1.1.529, BA) variant within US surveillance networks. Several tracked Omicron (BA) lineages (*e*.*g*., BA.2.75, BA.2.75.2, BF.11, and BA.5.2.6) were detected at low levels (≤10%, labels not shown). “Prior lineages” included additional A and B lineages with limited detection. “Other lineages” consolidated the few specimens with no assigned Pango nomenclature or non-XBB X lineage.

The U.S. CDC’s national SARS-CoV-2 surveillance system collects SARS-CoV-2 specimens for sequencing through the National SARS-CoV-2 Strain Surveillance (NS3) program and various laboratories. Besides the predominant Omicron lineages, other less prevalent lineages were also identified during the surveillance. As part of CDC’s effort to assess the risk of emerging SARS-CoV-2 variants (primarily Omicron lineages in 2022), we continuously assessed the antibody neutralization of viruses representative of emerging Omicron lineages. Selection of viruses lineages to test was based on epidemiologic data, lineage proportion, forecasted growth, spike mutations, and other information reported by public health and academic laboratories. Sera collected from volunteer vaccinees who received three doses of the original monovalent mRNA vaccine (based on the index virus) were used in neutralization assays throughout 2022. Sera from vaccinees who received a fourth dose of the bivalent mRNA vaccine became available in late 2022, and those sera were also used in assays of lineages that emerged more recently.

In this study, we assessed the neutralizing activity of sera from post-third monovalent dose and post-fourth bivalent dose against 10 Omicron lineages (i.e., BA.1, BA.2, BA.5, BA.2.75, BA.2.75.2, BN.1, BQ.1, BQ.1.1, XBB, and XBB.1), which included lineages that predominated in different periods of 2022 as well as recent co-circulating lineages with the most significant changes in the spike. We further dissected the role that the receptor-binding domain (RBD) and non-RBD spike mutations played in the observed antibody escape by uncoupling mutations identified in lineages with different spike proteins.

## Results

### Anti-spike and anti-RBD antibody binding activity of post-third (monovalent) dose and post-fourth (bivalent) dose sera

Sera were collected from vaccinees 2-6 weeks after receiving a third dose of monovalent mRNA vaccine or 2-7 weeks after receiving a fourth dose of bivalent booster mRNA vaccine. Meso Scale Discovery (MSD) assays were used to compare the total IgG antibody response (i.e., neutralizing and non-neutralizing) elicited by post-third dose sera and the post-fourth dose sera against representative spike proteins of the progenitor index virus (614D), Alpha, Beta, Delta, and Omicron lineages of BA.1, BA.2, BA.5, and BA.2.75. Anti-spike binding activity post-third dose showed minimal decrease for Alpha, Beta, and Delta, with fold reductions of 1.2, 1.7, and 1.4 (350,666 AU/mL, 257,039 AU/mL, and 293,492 AU/mL) compared to the 614D reference (425,339 AU/mL), respectively. Larger differences were observed for BA.1, BA.2, BA.5, and BA.2.75, with fold-change decreases of 4.2, 3.8, 3.7, and 5.6 (100,556 AU/mL, 112,390 AU/mL, 113,795 AU/mL, and 75,406 AU/mL), respectively (Fig. 2a). The post-bivalent vaccine (*i*.*e*., fourth dose) sera, compared to the post-third dose sera, showed slightly higher activity against all the spike antigens (Fig. 2b). However, the breadth/specificity of those two sera were similar in the MSD assay, as the fold changes compared to the 614D spike reference amongst BA.1, BA.2, BA.5, and BA.2.75 between post-third dose and post-bivalent sera remained similar (i.e., 4.2 vs. 3.8, 3.8 vs. 3.1, 3.7 vs. 2.9, and 5.6 vs. 4.9) (Fig. 2a and 2b). When the RBD domain was used as the target antigen, the antibody binding decreased an additional ∼3-fold for both sera against all variants and Omicron lineages (Fig. 2c and 2d). However, again, the fold changes amongst BA.1, BA.2, BA.5, and BA.2.75 between post-third dose and post-bivalent sera remained similar (4.6 vs. 4.4, 3.6 vs. 3.4, 4.0 vs. 4.1, and 4.8 vs. 4.7) compared to the 614D RBD reference (Fig. 2c and 2d).

**Fig. 2:**
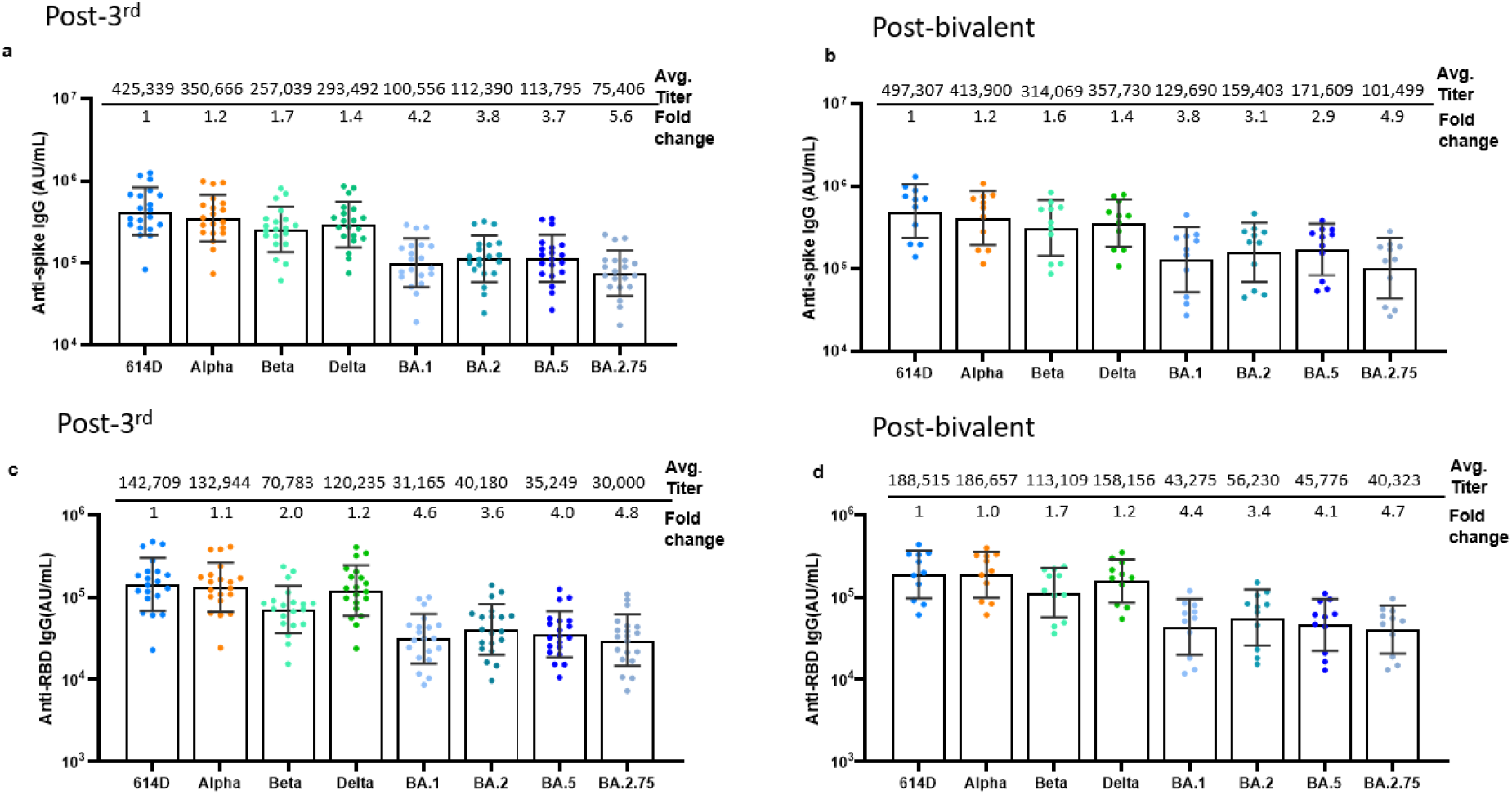
Quantification of anti-spike and anti-RBD IgG binding activity in post-third dose and post-fourth dose (bivalent) vaccinee sera. Each dot represents the IgG measured in AU/mL for one serum-antigen pair. Geometric mean titers with geometric standard deviations are displayed across the top of each graph. Fold-change was calculated as the ratios relative to 614D. For statistical analysis, a two-tailed Wilcoxon matched-pairs signed-rank test was performed by comparing each variant with 614D. Test statistics and P value are summarized in Supplementary Table 2. IgG antibodies were measured for 8 different spikes and RBDs as: **a**, anti-spike antibodies in post-third dose sera (N = 20 biologically independent post-third dose sera examined over 8 spike antigens); **b**, anti-spike antibodies in post-bivalent dose sera (N = 11 biologically independent post-bivalent dose sera examined over 8 spike antigens); **c**, anti-RBD antibodies in post-third dose sera (N = 20 biologically independent third-dose sera examined over 8 RBD antigens); and **d**, anti-RBD antibodies in post-bivalent dose sera (N = 11 biologically independent post-bivalent dose sera examined over 8 RBD antigens). Variant and lineage names are displayed across the bottom of each graph. Within each serum-antigen pair, average IgG concentrations were significantly lower (P < 0.001) for all viruses tested relative to the 614D reference except for Alpha in Fig. 2d (P = 0.8984). AU, arbitrary units.

It will be interesting to compare the post-third dose and post-bivalent sera against the spike or RBD of more recent Omicron lineages, such as BQ.1.1 and XBB.1, once the corresponding MSD kits are manufactured. However, based on the less-than-2-fold difference against the different variants analyzed (Fig. 2), the spike-binding antibody level against these more recent Omicron lineages in the post-third dose sera and post-bivalent is likely to be within 2-fold.

### Neutralizing activity of post-third dose and post-fourth (bivalent) dose sera against Omicron lineages

To determine the neutralizing activity of post-third dose and post-bivalent vaccine dose sera against key Omicron lineage viruses, we generated a recombinant BA.1 fluorescent reporter virus and engineered lineage-specific spike mutations in the BA.1 spike background, resulting in corresponding reporter viruses for 10 Omicron lineage viruses (Supplementary Fig. 1). These reporter viruses contained all the spike and non-spike genes (minus ORF7) from Omicron and an mNeonGreen fluorescent reporter gene in place of ORF7 to enable rapid high-throughput focus reduction neutralization test (FRNT) in the context of SARS-CoV-2 rather than pseudotyped viruses. BA.1, BA.2, and BA.5 represent the three predominant Omicron lineages in 2022 from which other lineages have evolved (Fig. 1). BA.2 spawned the BA.2.75 lineage which evolved into BA.2.75.2 and BN.1. BA.2.75 underwent a recombination event with BJ.1 (a descendant of BA.2.10) to form the XBB and XBB.1 recombinant lineages^9^, which predominated in Singapore^10^ and are also increasing in the U.S. BA.5 was also derived from BA.2, and its descendant lineages, BQ.1 and BQ.1.1, were the predominant lineages in the U.S. in November and December 2022. While the evolution of SARS-CoV-2 RNA virus genome is complex, the spike proteins encoded by these Omicron lineage viruses are fairly closely related with the XBB.1 virus being the most divergent (Supplementary Fig. 1).

The post-third dose sera showed high neutralizing activity against the 614D reference virus (Neutralizing antibody (nAb) titer = 3,451 IU/mL), intermediate activity against the BA.1, BA.2, BA.5, and BA.2.75 (nAb titer = 174 – 647 IU/mL), and low activity against BA.2.75.2, BN.1, BQ.1, BQ.1.1, XBB, and XBB.1 (nAb titer = 9 – 54 IU/mL) (Fig. 3a). Compared to the nAb titer of 614D, the nAb titers of BA.1, BA.2, BA.5, and BA.2.75 decreased by 5-, 8-, 20- and 12-fold, respectively. Titers of the two BA.2.75-derived lineages, BA.2.75-2 and BN.1, decreased by 127- and 64-fold compared to 614D, respectively. Titers of the BA.5-derived lineages, BQ.1 and BQ.1.1, decreased by 190- and 288-fold compared to 614D, respectively. XBB and XBB.1 exhibited the greatest extent of escape, with 324-fold and 371-fold nAb reductions compared to 614D, respectively (Fig. 3a).

**Fig. 3:**
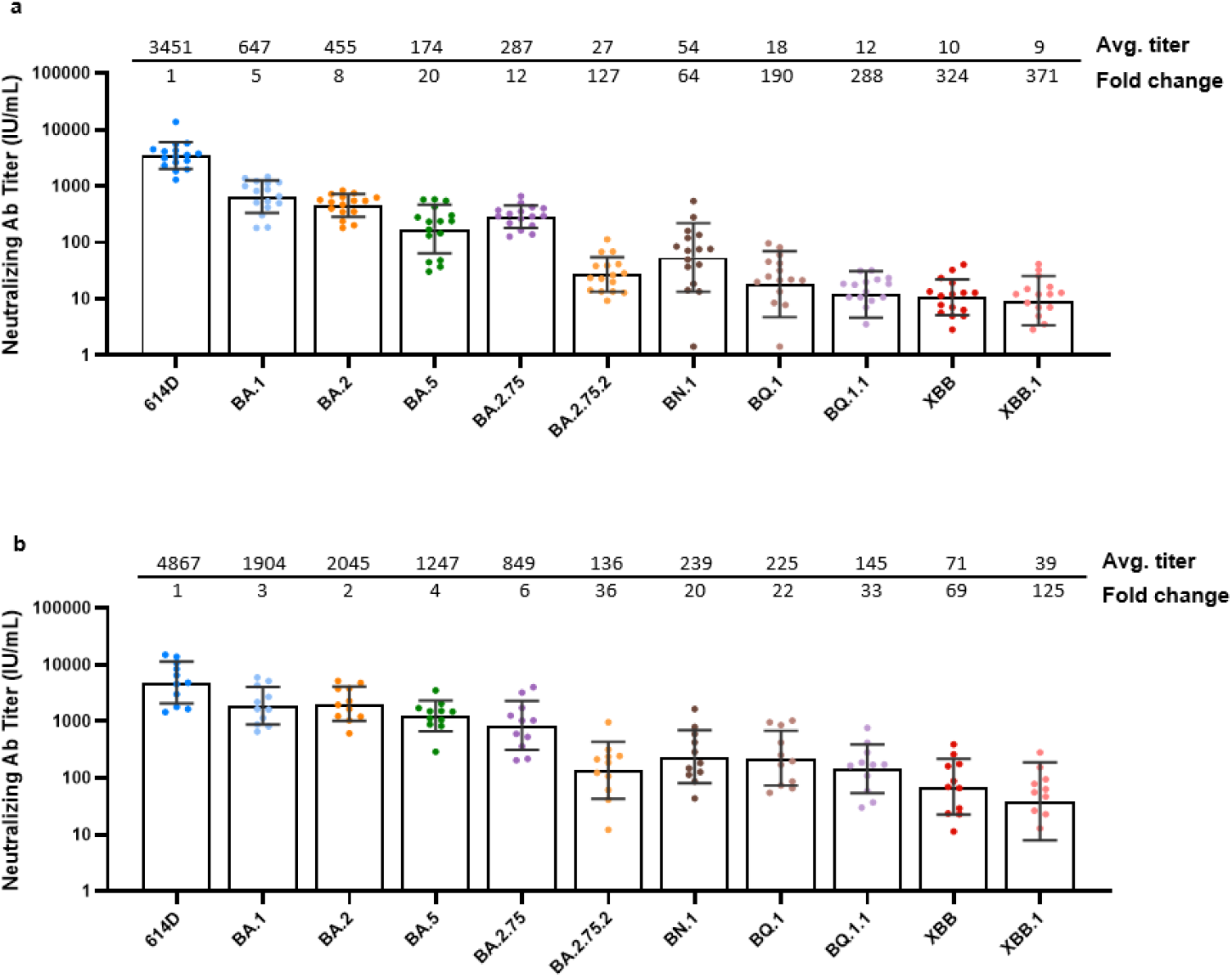
Neutralization activity of post-third (monovalent) dose sera and post-fourth (bivalent) dose sera against Omicron lineages. Each dot represents the neutralizing antibody titer (IU/mL) for each serum-virus pair. For statistical analysis, a two-tailed Wilcoxon matched-pairs signed-rank test was performed by comparing each variant with 614D. Test statistics and P value are summarized in Supplementary Table 2. The geometric mean of neutralizing antibody titer with geometric standard deviation and average fold-change with respect to the 614D reference virus are displayed across the top of each graph for: **a**, post-third dose sera (8 Pfizer-BioNTech BNT162b2 mRNA vaccine and 8 Moderna mRNA-1273 vaccine biologically independent post-third dose sera examined over 11 viruses); and **b**, post-fourth dose bivalent vaccine sera (N = 11 biologically independent post-fourth dose bivalent vaccine sera examined over 11 viruses). Average fold-change was calculated as the individual ratios of the geometric mean relative to 614D. Average neutralizing antibody titers in both sera for all lineages differ significantly (P < 0.001) from 614D (Supplementary Table 2). Virus lineage names are displayed across the bottom of each graph.

Compared to the post-third dose sera, neutralizing activity against the 614D reference increased moderately, by 1.4-fold, with the post-bivalent vaccine sera (nAb titer = 4,867 vs. 3,451 IU/mL), but dramatically increased against all Omicron lineages (Fig. 3b). Specifically, the post-bivalent vaccine sera were 7.1-fold more potent in neutralizing BA.5 (homologous antigen) and 2.9- to 5.1-fold more potent in neutralizing heterologous antigens BA.1, BA.2, BA.2.75, BA.2.75.2 and BN.1, than the post-third dose sera (Fig. 3a vs. Fig. 3b). The greater increase of BA.5 compared to BA.1/BA.2 lineage viruses indicated some BA.5 lineage-specific antibodies were elicited by the bivalent mRNA vaccine. This was corroborated by the observation that the post-bivalent vaccine sera were 12-fold more potent than the post-third dose sera in neutralizing the currently predominant BQ.1 (nAb titer = 225 vs. 18 IU/mL) and BQ.1.1 (nAb titer = 145 vs. 12 IU/mL), which descended from BA.5 and have nearly identical spike glycoproteins (Fig. 3b and Supplemental Fig. 1).

In summary, these results demonstrated that the bivalent booster vaccine that included the Omicron BA.4/BA.5 antigen increased neutralizing antibodies against progenitor and contemporary Omicron lineages, improved antibody breadth against all Omicron lineages, and elicited greater effect on BA.5 lineage viruses (*e*.*g*., BA.5, BQ.1, and BQ.1.1) than on other Omicron lineages (*e*.*g*., BA.1, BA.2.75.2, XBB).

### Impact of RBD and non-RBD mutations on neutralizing activity

There are numerous mutations in the spike glycoprotein of Omicron lineages compared to the index virus and the B.1.1.529/BA.1 virus represented a major antigenic drift that led to a global sweep. More mutations accumulated in the spike during the evolution of the Omicron in human population. For example, the XBB.1 lineage carries approximately 40 spike glycoprotein mutations, roughly half of which reside within the RBD, which contains epitopes targeted by potent neutralizing antibodies (Supplementary Fig. 1). It is important to understand the molecular mechanism (e.g., domains/specific mutations) responsible for the most significant neutralization escape because these data inform risk assessment and/or vaccine antigen selection or design. To specifically analyze the role of different spike domains playing in antibody escape, we chose the lineages BA.1, BA.2, BA.5, and XBB.1 and segregated their RBD and non-RBD spike mutations. All the viruses were analyzed using pooled post-second dose sera (primary series of two doses of mRNA vaccine) and pooled post-third dose sera. The post-second dose sera were included to accurately measure the reduction in neutralization for SARS-CoV-2 viruses containing different spike domains, as we found the post-second dose sera were more sensitive to antibody escape than the post-third dose sera^2^.

Compared to the 614D virus, titer reductions for the BA.1, BA.2, BA.5, and XBB.1 were 45-, 38-, 29- and >192-fold for post-second dose, respectively, and 8-, 11-, 10- and 289-fold for post-third dose, respectively (Supplementary Table 1). Viruses carrying only RBD mutations (*i*.*e*., BA.1-RBD, BA.2-RBD, BA.5-RBD, and XBB.1-RBD) were associated with titer reductions of 1-, 3-, 7- and 38-fold for post-second dose, respectively, and 1-, 4-, 8- and 17-fold for post-third dose, respectively. Removing the RBD mutations (*i*.*e*., BA.1-wt-RBD, BA.2-wt-RBD, BA.5-wt-RBD, and XBB.1-wt-RBD) restored neutralizing titers of the viruses to near 614D levels (1- to 2-fold difference) in both post-second and post-third sera (Supplementary Table 1). Because neutralizing titers against the XBB.1 virus lacking RBD mutations (XBB.1-wt-RBD) were comparable to those of 614D (1- to 2-fold reduction), the 38-fold (post-second dose) and 17-fold (post-third dose) reductions obtained using the XBB.1-RBD virus cannot explain the >192-fold (post-second dose) and 289-fold (post-third dose) reductions of XBB.1 with full spike mutations. Previous studies using a domain-specific antibody-depletion strategy have shown that the RBD accounts for up to 90% of the targeted neutralization activity^11 12, 13, 14^. Our strategy of using live SARS-CoV-2 viruses with Omicron spike mutations removed from different regions suggests that spike mutations outside RBD may function synergistically with RBD mutations resulting in a dramatic escape from antibody neutralization by XBB.1. This underscores the importance of mutations outside the RBD region in the extreme antibody escape by XBB.1, probably by inducing conformational changes in the spike structure or by increasing fitness in other ways such as higher stability, tighter receptor binding, and more efficient fusion.

## Discussion

Two recent CDC-led vaccine effectiveness (VE) studies found that the bivalent mRNA vaccine booster provided 42-73% additional protection against COVID-19-associated hospitalization compared with 2 to 4 monovalent COVID-19 vaccine doses during September-November 2022^15, 16^, when BA.5 and BQ.1/BQ.1.1 predominated. In the present study, from the post-third dose monovalent sera to post-fourth dose bivalent vaccine sera, the neutralizing titers increased by 7.1-fold against BA.5 and 12-fold against BQ.1 and BQ.1.1 (Fig. 3), strongly suggesting that the higher VE was resulted from the increased neutralizing titers from the bivalent vaccine. While the correlates of protection to SARS-CoV-2 continue remain to be fully elucidated, they may be further established as we collect more data from the VE studies and determine serum neutralizing titers of corresponding vaccinees in the following months.

We recognize a limitation of this study may be the lack of fourth dose monovalent sera as a comparison. We have previously shown that neutralization activity increased following a third dose of monovalent booster compared to the primary series of two vaccine doses^2^. The boosted activity not only increased against 614D by 4-fold but also against Omicron lineages BA.1 and BA.2 by 13-fold and 9-fold, respectively, suggesting increased breadth^2^. However, a more recent study on Omicron lineages showed minimal effect on antibody breadth comparing post-third dose monovalent and post-fourth dose monovalent vaccination, where neutralizing titers increased evenly for the D614G virus (2.8-fold), BA.2 (2.7-fold), BA.4/5 (2.5-fold), and BQ.1 (2.4-fold)^17^. Thus, additional monovalent doses of the original mRNA vaccine alone may not consistently increase the breadth of protection. Lineage-specific boosters and/or antigens designed to enhance breadth are likely essential to combat future antigenic changes.

Recent studies to define SARS-CoV-2 immune correlates of protection have related an increase of neutralizing titers from 10 to 100 IU/mL with respective increases in VE from 78% to 91%^18^. Additional studies have reported neutralization titers increasing from 8 IU/ml to 26 IU/ml in association with VE increasing from 70% to 80%^19^ and from 9.9 IU_50_/ml to 96.3 IU/ml with VE from 78% to 89%^20^. In the present study, although the post-third dose sera only had neutralizing titers of ∼10 IU/mL against BQ.1.1, XBB, and XBB.1, the post-bivalent vaccine sera increased the titers to 145 IU/mL for BQ.1.1, 71 IU/mL for XBB, and 39 IU/mL for XBB.1, a similar trend as reported recently by others^21, 22, 23, 24^. Sera used in this test were collected 2-7 weeks post bivalent booster, and considering the waning of antibodies over time of 2- to 10-fold decreases in about 6 months^25, 26, 27, 28, 29^, the bivalent vaccine booster may protect against these lineages for a few more months but unlikely for more than 6 months. Furthermore, the effect of waning immunity increases as the virus spike mutates and the low titers of XBB variants illustrates these and their descendants are most likely to escape vaccine and infection induced immunity resulting in symptomatic infections. Nevertheless, the increased level of polyclonal antibodies induced by the bivalent vaccine (Fig. 2) should reduce virus replication through opsonization and/or antibody dependent cellular cytotoxicity, aiding in protection from severe disease. Furthermore, cell mediated immunity is largely unaffected by the relatively small number of mutations in the spike or other parts of the genome^30, 31^. Another booster with bivalent, or monovalent BA.4/BA.5 vaccine will likely help, but continual updates to vaccine antigens to induce antibodies that better protect against circulating and future lineages are likely to offer superior protection.

Collectively, rapid evolution of Omicron lineages warrants continued close epidemiologic and virologic surveillance that includes continual virus neutralization analysis of emerging viruses. Vaccination remains the most effective strategy to combat the COVID-19 pandemic.

## Methods

### Ethics statement

Vaccinee serum samples were collected from individuals through the Investigating Respiratory Viruses in the Acutely Ill (IVY) Network, a Centers for Disease Control and Prevention (CDC)-funded collaboration to monitor the effectiveness of SARS-CoV-2 vaccines among US adults. Participants were vaccinated with a primary series (initial two doses) of either the Moderna mRNA-1273 or Pfizer-BioNTech BNT162b2 vaccine, and then boosted with either a third monovalent dose or, additionally, a fourth bivalent dose. Post-second dose sera^2^, post-third dose sera^2^, and post-fourth dose sera were collected 2-6 weeks, 2-6 weeks or 2-7 weeks after vaccination, respectively. Participants had no prior diagnosis of infection with SARS-CoV-2 based on self-reporting or low levels of anti-nucleocapsid protein antibodies in MSD assay. This activity was approved by each participating institution, either as a research project with written informed consent or as a public health surveillance project without written informed consent. This activity was also reviewed by the CDC and conducted in a manner consistent with applicable federal laws and CDC policies: see e.g., 45 C.F.R. part 46.102(l)(2), 21 C.F.R. part 56; 42 U.S.C. §241(d); 5 U.S.C. §552a; 44 U.S.C. §3501 et seq.

### Biosafety statement

Infectious SARS-CoV-2 viral work was performed under enhanced BSL-3E conditions. Additional safety measures were followed as described^2^.

### SARS-CoV-2 variant and Omicron lineage prevalence

SARS-CoV-2 variant and lineage analytics for NCBI- and GISAID-reported specimens within the National SARS-CoV-2 Strain Surveillance (NS3) network, CDC-contracted diagnostic and research laboratories, and baseline US surveillance were rendered in Tableau Desktop (version 2022.3.0). Daily, percent proportionalities were aggregated by attributed Pango annotation (version 4.1.3) with further consolidation based on B.1.1.529 (BA) lineages: BA.1, BA.1.1, BA.2, BA.2.12.1, BA.2.75, BA.2.75.2, BA.3, BA.4, BA.4.6, BA.5, and BA.5.2.6. In addition, select BA.2.75.5 (BN; BN.1), BA.5.2.1 (BF; BF.7 and BF.11), and BA.5.3.1.1.1.1.1 (BQ; BQ.1, and BQ.1.1) lineages were included. Exceptions included the non-BA lineages: B.1.617.2 (Delta) and XBB. All A and remaining B lineages were assigned the “Prior Lineage(s)” label. Both non-XBB X lineages and specimens without an assigned Pango nomenclature were consolidated into “Other Lineage(s).” Applied analytics included available surveillance data from November 1, 2021, to November 30, 2022 (last updated on December 19, 2022).

### Generating SARS-CoV-2 reporter viruses

RNA was extracted from a SARS-CoV-2 Omicron BA.1 clinical isolate (SARS-CoV-2/human/USA/CA-CDC-4358237-001/2021) (GenBank: OM264909.1) using the QIAamp Viral RNA kit (Qiagen, Hilden, Germany). The complete wild-type Omicron genome was then reverse transcribed via RT-PCR with SuperScript™ IV First-Strand Synthesis System (Thermo Fisher Scientific, Waltham, MA, 18091050), amplified with Q5 High-Fidelity DNA Polymerase (NEB, Ipswich, MA, M0492S), and cloned into a bacterial artificial chromosome as described^2^. Reverse genetics SARS-CoV-2 clones were produced as previously described^2^ with a mNeonGreen fluorescent reporter gene^32, 33^. *In vitro* transcribed RNA from these infectious clones was then electroporated into VeroE6-N cells as described^2^. Sequence of rescued viruses was confirmed by Illumina Next-generation sequencing. Domain substitution analysis used the SARS-CoV-2 index virus as reference.

### Meso scale discovery (MSD) immunoassay

Meso Scale Discovery (MSD) assays were performed as previously described^2^. Briefly, 5 μl of each post-third dose and post-bivalent vaccine dose serum was added to 245 μl of buffer, initially diluted by 1:50, and then serially diluted by 1:10. The 1:500 dilution was used to measure IgG binding to SARS-CoV-2 nucleocapsid protein to exclude sera with prior infection. The final 1:50,000 dilution was used to measure IgG binding to SARS-CoV-2 spike from a variety of variants and Omicron lineages, including 614D, Alpha, Beta, Delta, BA.1, BA.2, BA.5, and BA.2.75, in addition to the RBD alone. Binding was measured using the V-PLEX SARS-CoV-2 Key Variant Spike Panel 1 Kit (K15651U-2 (IgG)) and Key Variant RBD Panel 1 Kit (K15659U-2 (IgG)). IgG concentrations were calculated using the DISCOVERY WORKBENCH 4.0 Analysis Software.

### Focus reduction neutralization test (FRNT)

FRNT was used to determine serum neutralization titers and performed as previously described^2^. Briefly, post-third dose and post-bivalent vaccine sera were serially diluted in 3-fold steps for 7 dilutions (starting from 1:10 or 1:30) in sextuplicate in 96-well round bottom plates. SARS-CoV-2 reporter virus was diluted to 3,200-6,000 focus forming units (FFUs) per mL. Diluted serum samples were mixed with an equal volume of diluted virus and incubated for 1 hour at room temperature (21±2°C). Serum-virus mixtures were then inoculated into each well of VeroE6/TMPRSS2 cells in 96-well tissue culture plates and incubated at 37°C in a 5% CO_2_ atmosphere for 2 hours. The wells were then overlaid with 100 μl of 0.75% methylcellulose and incubated at 33°C in a 5% CO_2_ incubator for 16-18 hours. Plates were scanned on the Cytation7 and FFU were quantified using Gen5 version 3.11 (BioTek). FRNT_50_ values were calculated from FFU values as previously described^2^. The First WHO International Standard for anti-SARS-CoV-2 immunoglobulin (human; NIBSC code: 20/136) was included in the assay to calibrate the neutralizing antibody titer in International Units per milliliter (IU/mL)^34^.

### Statistical analysis

Average fold changes were calculated using the geometric mean of neutralizing Ab titer ratios against that of 614D. Statistical analyses were performed using GraphPad Prism 9.3.1 and R version 4.1.2, with significance defined as P < 0.05. A two-tailed Wilcoxon matched-pairs signed-rank test was used to determine significance of anti-spike and anti-RBD IgG titers as well as neutralizing antibody titers relative to 614D (Supplementary Table 2).

## Data availability

SARS-CoV-2 genome sequences of viruses generated in this study are being deposited in GenBank. Additional data generated from this study are provided in the Supplementary Information/Source Data files.

## Code availability

Scripts used to calculate the FRNT_50_ titers were deposited previously^2^.

## Acknowledgements

We thank the Strain Surveillance and Emerging Variant Team’s Bioinformatics Unit for their work in tracking the evolution of the US SARS-CoV-2 viruses: Peter Cook, Adam Retchless, Sandra Mathew, Philip Shirk Catherine Smith, Xiao-Yu Zheng, Samuel Shepard, Thomas Stark, Dakota Howard, Roopa Nagilla, Nicole Paterson, David Patton, Jason Caravas, and Dhwani Batra. We acknowledge the investigators in the SARS-CoV-2 Assessment of Viral Evolution (SAVE) Program under the National Institute of Allergy and Infectious Diseases (NIAID) for tracking and sharing of information on spike mutations in emerging SARS-CoV-2 variants and lineages. This study was funded by the CDC COVID-19 Emergency Response. The findings and conclusions in this report are those of the authors and do not necessarily represent the official position of the U.S. Centers for Disease Control and Prevention or the Agency for Toxic Substances and Disease Registry.

## Author contributions

D.E.W. and B.Z. conceived the study. L.W. and B.Z. designed the experiments. N.J., LW., D.E.W., and B.Z. wrote the manuscript. N.J., L.W., M.H., C.F., M.C., X.L., J.H., D.C., B.R.M., W.W., G.A., M.W., R.C., B.R., J.R.B, and B.Z. performed experiments and/or analyzed data. B.R.M., N.K., K.L., C.R.P., N.H., R.J.K, and A.S.L. tracked and analyzed the prevalence of SARS-CoV-2 variants and Omicron lineages. W.H.S., J.P.R., A.B., J.D.C., N.I.S., K.W.G., D.N.H., D.S., M.L.M., and N.J.T. supervised or coordinated serum sample collection. All authors have reviewed and approved this manuscript.

## Figures

**Supplementary Fig. 1:**
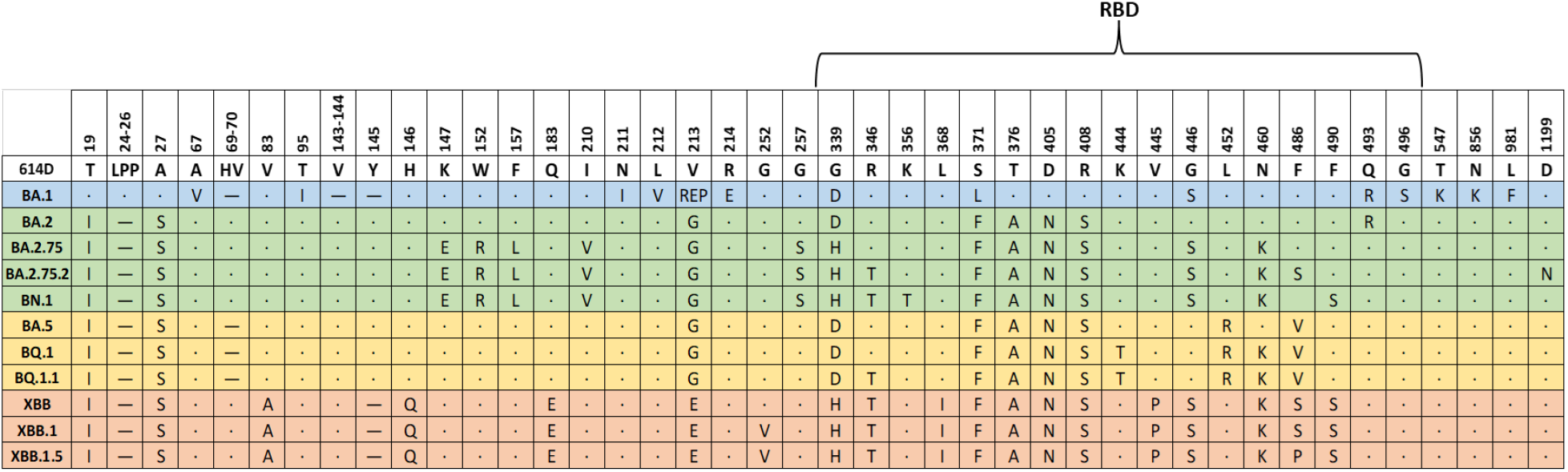
Omicron lineage spike mutations relative to 614D. Spike mutations in Omicron lineages BA.1, BA.2, BA.2.75, BA.2.75.2, BN.1, BA.5, BQ.1, BQ.1.1, XBB, XBB.1, and XBB.1.5 relative to 614D are shown. Mutations common across all 11 Omicron lineages are not displayed. Spike amino acid positions are displayed across the top of the figure with the RBD annotated. BA.1 is highlighted in blue. BA.2 and descendants are highlighted in green. BA.5 and descendants are highlighted in yellow. XBB and descendants are highlighted in orange. Dots represent no change relative to 614D. Dashes represent deletions.

